## Supplementary Information for "Tardigrade secretory proteins protect biological structures from desiccation"

**Table S1.** Information of SAHS proteins used in this study.

| Name | Uniprot ID | Host organism | Amino acid lengths (including secretion tag) |
|---|---|---|---|
| RvSAHS1 | J7MFT5 | *R. varieornatus* | 169 |
| RvSAHS2 | J7MAN2 | *R. varieornatus* | 174 |
| RvSAHS3 | A0A1D1UKM2 | *R. varieornatus* | 146 |
| RvSAHS4 | A0A1D1UN89 | *R. varieornatus* | 171 |
| HeSAHS4 | P0CU42 | *H. dujardini* | 174 |
| RvSAHS6 | A0A1D1UJP0 | *R. varieornatus* | 172 |
| RvSAHS7 | A0A1D1UQJ5 | *R. varieornatus* | 159 |
| RvSAHS8 | A0A1D1UJQ1 | *R. varieornatus* | 173 |
| RvSAHS9 | A0A1D1UKG0 | *R. varieornatus* | 113 |
| RvSAHS10 | A0A1D1UJR2 | *R. varieornatus* | 123 |
| RvSAHS11 | A0A1D1URS8 | *R. varieornatus* | 171 |

| | | | |
|---|---|---|---|
| RvSAHS12 | A0A1D1UNQ6 | *R. varieornatus* | 110 |

**Table S2.** Estimation of protein localization using TargetP software.

| Protein name | Other | Secretory | Mitochondrial |
|---|---|---|---|
| RvSAHS1 | .0001 | .9998 | .0001 |
| RvSAHS2 | .0001 | .9999 | 0 |
| RvSAHS3 | .0004 | .9995 | .0001 |
| RvSAHS4 | .0045 | .9946 | .0009 |
| HeSAHS4 | .0001 | .9997 | .0002 |
| RvSAHS6 | .0009 | .9989 | .0003 |
| RvSAHS7 | .9989 | .0007 | .0004 |
| RvSAHS8 | 0 | 1 | 0 |
| RvSAHS9 | .9995 | .0003 | .0001 |
| RvSAHS10 | .9931 | .0004 | .0065 |
| RvSAHS11 | .0007 | .9977 | .0016 |
| RvSAHS12 | .9995 | .0002 | .0003 |

**Table S3. Amino acid sequences of the SAHS proteins (after SUMO cleavage).**

| Protein name | Amino acid sequence | Mw(kDa) |
|---|---|---|
| RvSAHS1 | APAEGHDDAKAEWTGKSWMGKWESTDRIENFDAFISALGLPLEQ<br>YGGNHKTFHKIWKEGDHYHHQISVPDKNYKNDVNFKLNEEGTTQ<br>HNNTEIKYKYTEDGGNLKAEVHVPSRNKVIHDEYKVNGDELEKT<br>YKVGDVTAKRWYKKSSSS | 17.3 |
| RvSAHS4 | RPHDESKAQWTGKPWLGKWESIDGTPENWEAFVKAANIPPKDQA<br>LYNGKQKTLLKYWKEAGEDHYHVQTSFPGTEHKMETSFKMGQEG<br>TLSHDGVDLKYVCTEDGEQLITKINIPSKNQETIVTYTATGDDL<br>EQTFTSNGVTGKRWYKKIHA | 17.3 |
| HeSAHS4 | TGDAPKEWSGKPWLGKFVAEVTDKSENWEAFVDALGLPEQFGRA<br>PVKTIQKIYKQGDHYHHIFALPDKNFEKDIEFTLGQEVEIKQGE<br>HIAKTKYSEDGEKLVADVSIPTKGKTIRSEYEVQGDQLIKTYKT<br>GDIVAKKWFKKVANPTEAPAQAA | 17.4 |
| RvSAHS6 | RPHDESKAQWTGKPWLGKWESTDKTPENWEAFVKAANIEPKYQS<br>LYSGKQKAIITIYKEGDSHYHAQMTFPGTDHKKEWDFKIGQEGT<br>YSMDGTEVKYVYTENGDQLDSKLNIPSKNTEMTHTYKVTGDELE<br>HIFTSNGATGKKWYKKVNNAV | 17.6 |

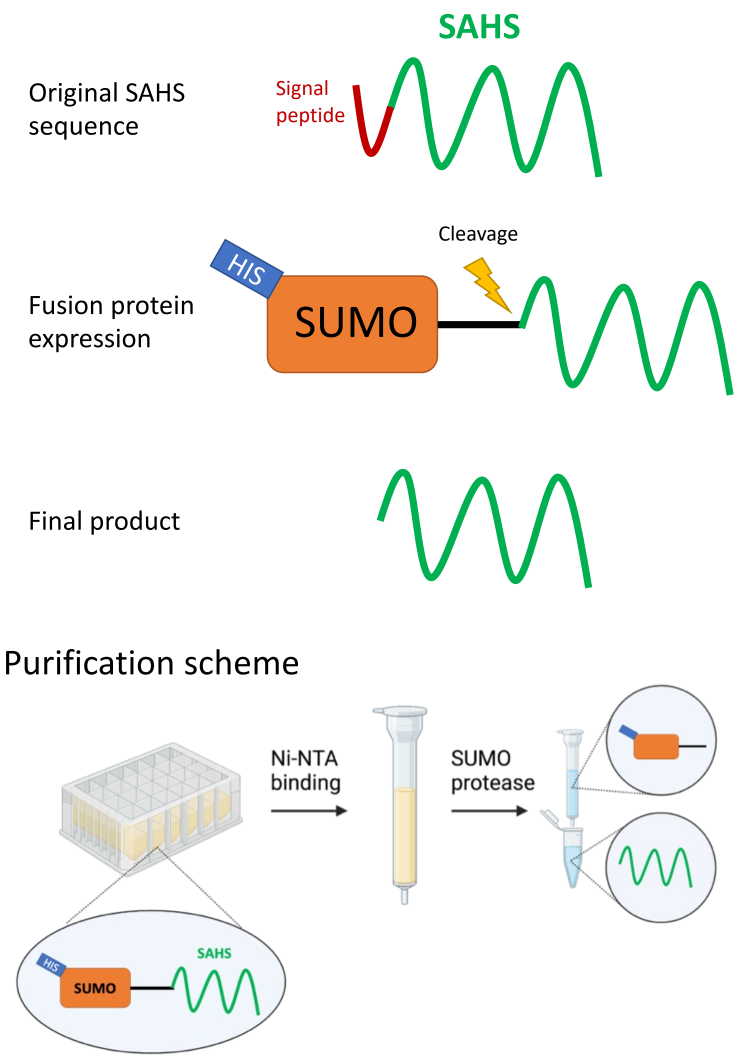


**Figure S1.** Sequence design and purification of the SAHS proteins.

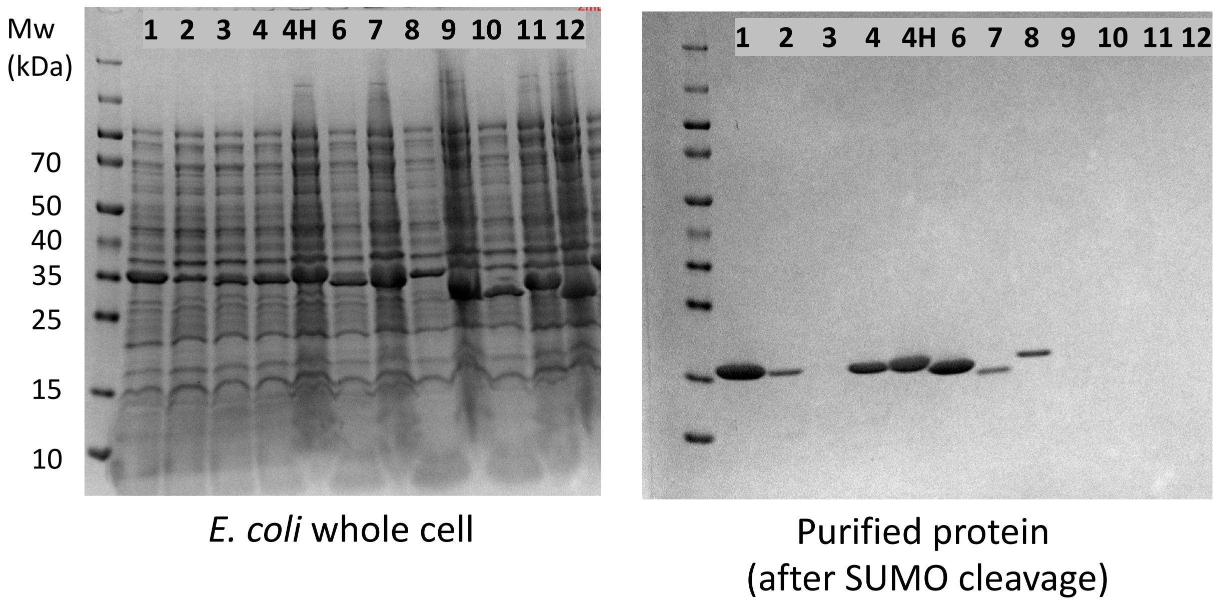


**Figure S2.** Expression and purification of the SAHS proteins from *E. Coli* host. Left gel shows the SDS-PAGE of *E. coli* whole cell expressing each SUMO-SAHS construct, and the right gel shows the final protein products after lysis, purification and SUMO cleavage. Each number indicates corresponding RvSAHS protein, and "4H" indicates HeSAHS4. RvSAHS1, 4, 6 and HeSAHS4 showed high soluble expression, whereas RvSAHS2, 7, 8 showed low soluble expression and the rest were expressed as insoluble fraction.

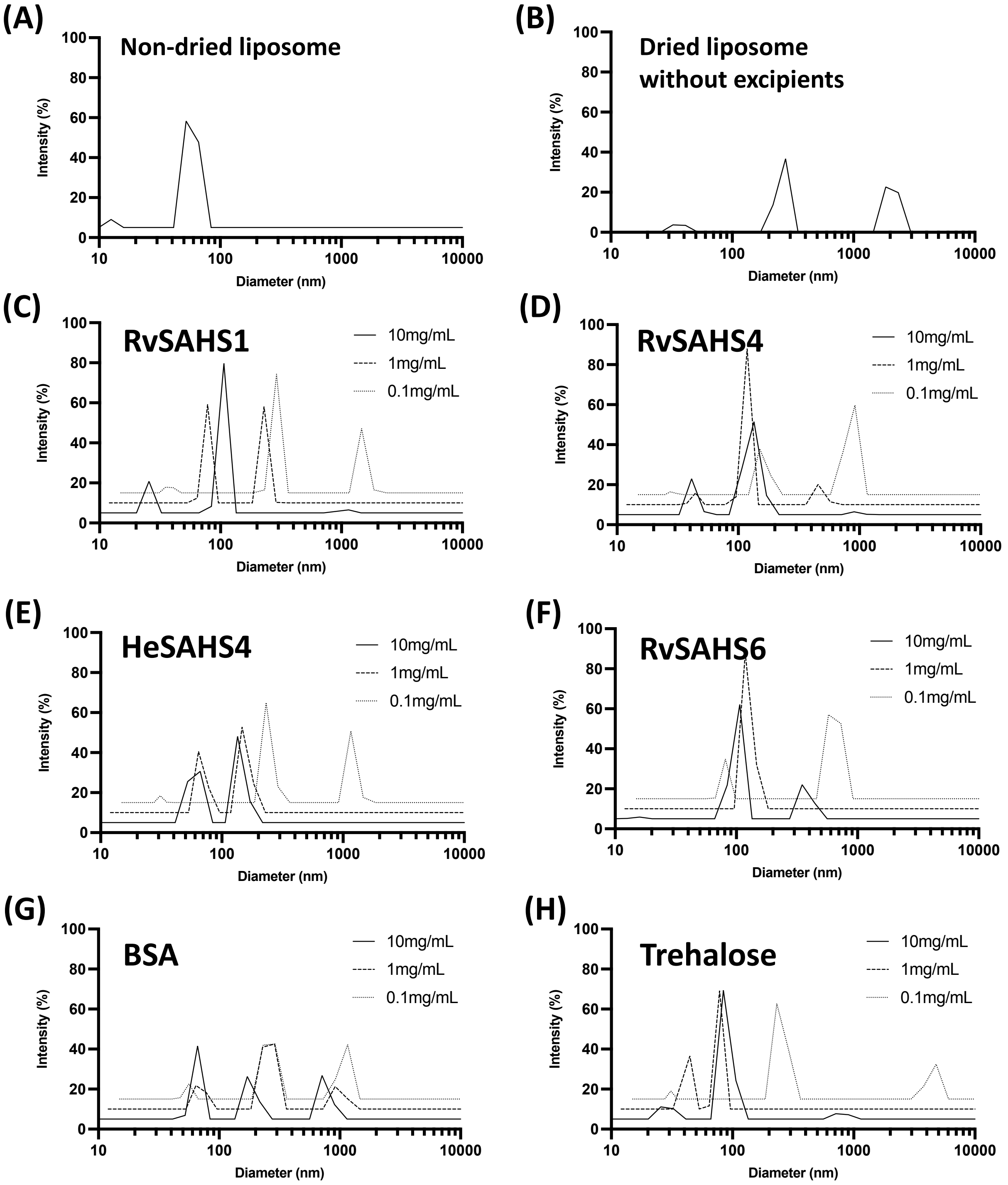


**Figure S3.** Data from a repeat experiment showing that SAHS proteins stabilize liposomes from desiccation-induced damages. POPC liposomes at 1.4 mg/mL were dried with and without the addition of SAHS proteins and BSA at varying concentrations of 0.1 – 10 mg/mL, and their size distributions were measured by DLS. (A) Size distribution of non-dried POPC liposomes (B) Size distribution of POPC liposomes dried and rehydrated without additives. (C-G) Size distributions of the liposomes dried with (C) RvSAHS1, (D) RvSAHS4, (E) HySAHS4, (F) RvSAHS6, (G) BSA and (H) trehalose.

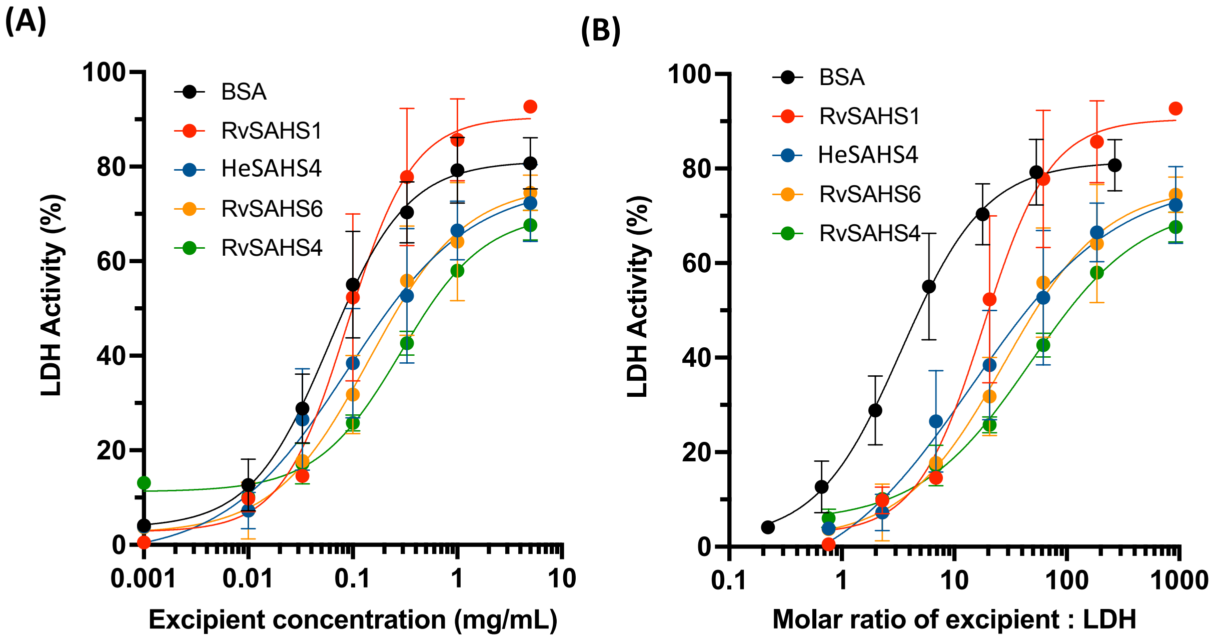


**Figure S4.** (A) Tardigrade SAHS proteins protect dried enzyme activity. LDH enzymes at 0.01 mg/mL were desiccated and rehydrated with and without the addition of SAHS proteins and BSA. Percent activity was determined using non-desiccated control samples stored at 4ºC as the reference to compare the activity. (B) The data from the same experiment as (A), depicted using molar ratio between excipient and LDH enzyme as the x-axis. Error bars represent the standard deviations from triplicate measurements.

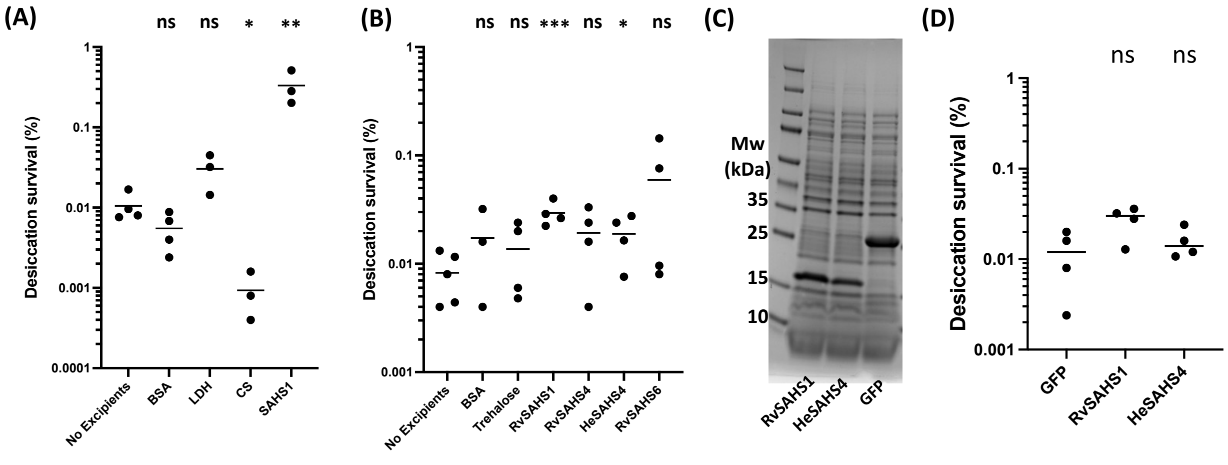


**Figure S5.** Desiccation protection of bacterial cells by SAHS proteins. (A) Desiccation survival percentage of *E. coli* cells dried for 48 hours with 0.5 mg/mL concentrations of control proteins including BSA, lactate dehydrogenase (LDH) and citrate synthase (CS). (B) Desiccation survival percentage of *E. coli* cells dried with 0.1 mg/mL concentration of extracellularly added SAHS proteins and control excipients. (C) SDS-PAGE of the whole *E. Coli* cells intracellularly overexpressing heterologous RvSAHS1, HySAHS4 and GFP. (D) Comparison of desiccation survival of the cells intracellularly expressing each protein. Individual data points represent independent replicates and lines represent the mean survival. Student's t-test was used to determine the statistical significance between the negative control (no excipient) and each group, which is indicated as asterisks. * $p < 0.05$; ** $p < 0.01$; *** $p < 0.001$.

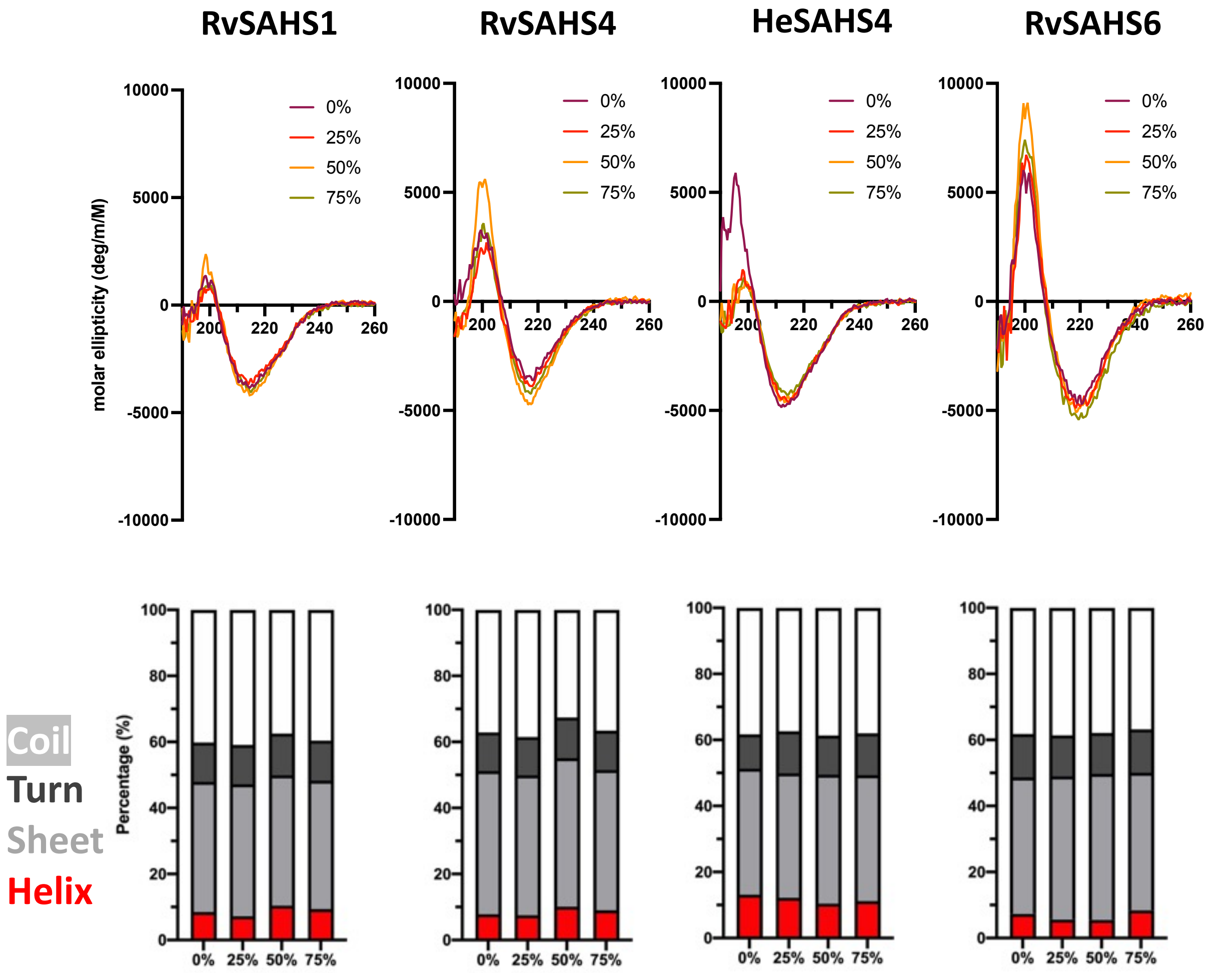


**Figure S6.** Effect of molecular crowding on SAHS protein structure. SAHS protein secondary structures upon glycerol addition were determined using circular dichroism. (A) CD spectra of SAHS proteins upon addition of increasing amounts of glycerol from 0 – 75%. (B) Secondary structure compositions of SAHS proteins under different glycerol level, calculated from the CD spectra.

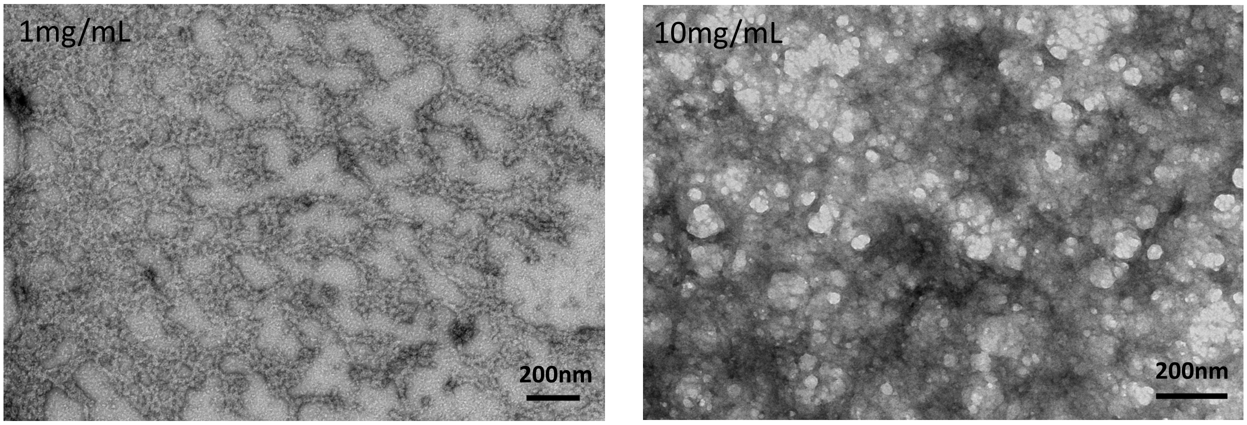


**Figure S7.** TEM images of the fibrous network structure formed by RvSAHS1 proteins dried at 1 mg/mL (left) and 10 mg/mL (right) concentrations. Scale bar = 200 nm.

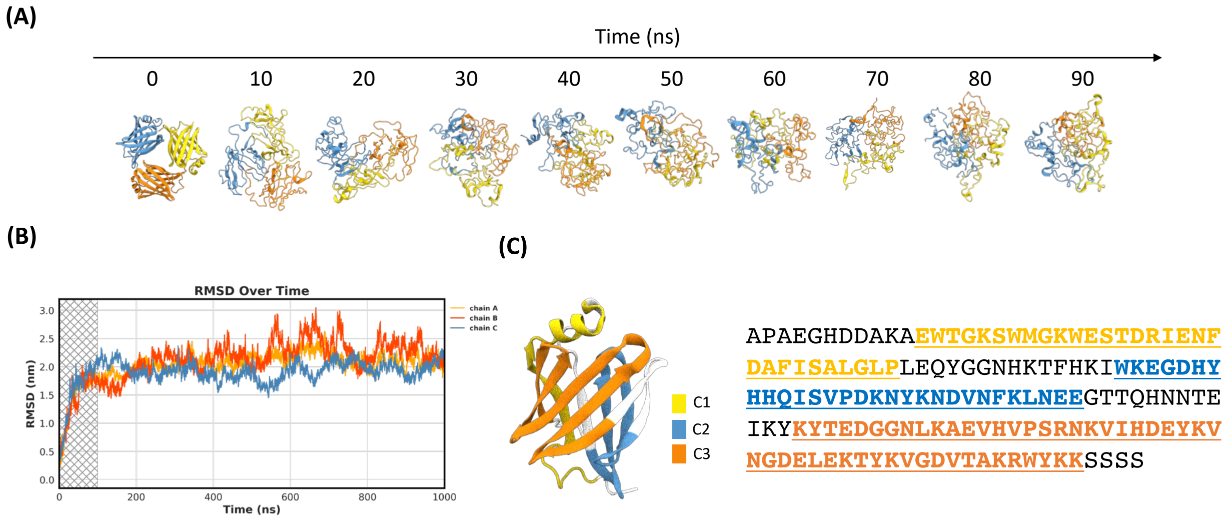


**Figure S7.** MD simulation of RvSAHS1 protein. (A) Changes in the structures of RvSAHS1 trimer during the first 100 ns of simulation. (B) RMSD over time during the entire 1 microsecond of MD simulation. Different color indicates each monomer chain in a trimer. (C) Conserved motifs C1-C3 of RvSAHS1 protein sequence in a single monomer. Amino acid sequence of RvSAHS1 is indicated, with C1 motif represented in yellow, C2 in blue, and C3 in orange, respectively.

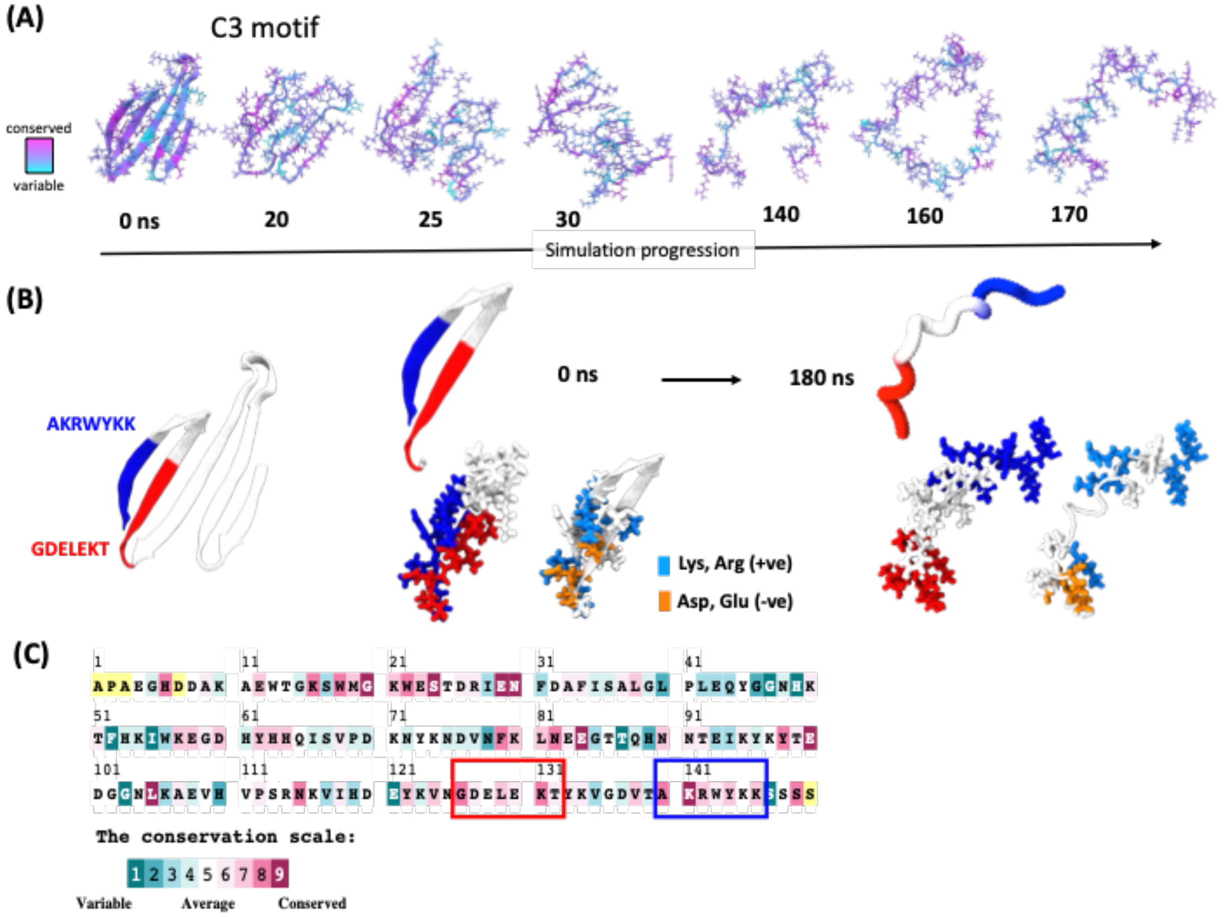


**Figure S8.** MD simulation showing structural changes in the conserved C3 motif of RvSAHS1. (A) Representative structural changes in C3 motif during the first 170 ns of simulation. Color scheme indicates the degree of evolutionary conservation, which shows that this motif is highly conserved among the SAHS family. (B) Close-up representation of the C3 motif sheet-to-helix structural change. Highlighted in blue and red are two highly conserved regions found within the beta sheet region that directly interact through ionic bonds. Structures of these regions at 0 and 180 ns are indicated, along with an additional depiction of the same regions in which positive residues (Lys, Arg) are highlighted in cyan and negative residues (Asp, Glu) are highlighted in orange. (C) Evolutionary conservation analysis of the RvSAHS1 sequence. Red and blue boxes indicate the same sequences depicted in (B).

**Supplementary Discussion**

The “short” SAHS proteins, SAHS9, 10 and 12 from *R. varieornatus* are truncated at their N-termini relative to other SAHS proteins. The Uniprot versions of SAHS9, 10 and 12 lack signal sequences, could not be expressed in a soluble form, lack residues that appear to be important in forming the hydrophobic core in SAHS1 and SAHS4, and for SAHS9 and SAHS12 began with a methionine corresponding to a methionine also found in SAHS3, 7 and 8. We therefore explored whether these might be incorrectly annotated and examined the genome sequence of *R. varieornatus* for upstream coding sequences that might have been overlooked. The results of this analysis are that the SAHS9, 10 and 12 genes encode proteins that may extend in the N-terminal direction further than the annotation would indicate, and that the new predicted protein sequences align well with N-termini of other SAHS proteins. However, we have not found clear-cut start codons for SAHS9 and 10, and SAHS12 may be much larger than the other SAHS proteins. The annotation below is intended to be a work in progress.

The SAHS9, 10 and 12 genomic regions were obtained from Genbank accession # BDGG01000001 (contig 1), a large contig from the *R. varieornatus* genome. [1] Relevant genome sequence fragments were obtained as reverse complements, then copy-pasted into Microsoft Word, and visually scanned for the presence of splice acceptors upstream of putative start codons for ORFs that translate into the Uniprot sequences. Because all of the SAHS genes in BDGG01000001 are on the anti-sense strand, we generated the reverse complement of this ~ 9Mb sequence for presentation purposes. The numbering used below is according to this reverse complement. Upstream regions of SAHS9, 10 and 12 were also copy-pasted into the Expasy Translate Tool (https://web.expasy.org/translate/), translated, and checked for long ORFs. A visual scan of the translated sequences revealed a pattern of tryptophans and other amino acids that align well with the N-terminus of other mature SAHS proteins (Figure 1B), but which are not found in the annotated sequences of these genes.

We also examined the intron/exon boundaries of these SAHS proteins and compared them to regions encoding other SAHS proteins to further validate the proposed mature amino acid sequences.

**SAHS9 analysis**

The SAHS9 protein sequence is most similar to SAHS10, 2 and 8. SAHS9 is also adjacent to SAHS2 and SAHS8 in the genome and is in a ~30 kilobase region that codes for SAHS11 plus SAHS10 (also adjacent) and SAHS7 plus SAHS1 (also adjacent).

When the SAHS9 genomic sequence is aligned with, for example that of SAHS2, there is a putative splice acceptor in SAHS9 DNA upstream of the annotated start codon at the same position as a splice acceptor annotated in SAHS2. Moving upstream, there is a putative splice donor in SAHS9 such that an intron of 79 bp is predicted, similar in size to the 82 bp intron in SAHS2. Moving further upstream, there is a predicted exon encoding a protein segment that aligns well with other full-length SAHS proteins, including Trp residues that help define the hydrophobic core. However, the predicted splice acceptor upstream of this exon has the sequence . . . TCCTTCTTACCG, lacking the canonical AG, and yet further upstream it is difficult to identify a splice donor and start codon that would align with other SAHS genomic sequences. It is possible that SAHS9 is a pseudogene, that this region of the sequence contains one or more errors, or that some other mechanism operates to express this gene.

Below is the genomic region encoding SAHS 9, 2 and 8.

```
 2575501 atttttcgcc tgtacagaca taagctgtta gagctgatgg agacacacct agatctcggg
 2575561 ttgacaccca atgacacggt aatgatcctt agaccattgt ccgaaaaatt gctcatcacc
 2575621 gttccctgta tggaaaaaaa ctggcttggc gggtaccttc gtgaacaata gtatgtcagt
```

2575681 tgagttttcg gtgagtcata aggggcagat agccgcgtgg cggaagcagc tgtttcggcc
2575741 tgtactgcaa accatgcata ggaaattggt ttaaaaaacg acttttccag ttttcttccc TATA box
2575801 tacatggata tttccgccca aatttagcca tccattcgtt tgtagaatgt ttccagcatt
2575861 cgataccita aggcatgctc ttgataacta cataagcggt ttcaagagtt cgactaacgc
2575921 ccctccagaa aacgtcgttt cctgaacagt ttagaaaata cctcggaacg gccaattaga splice donor
2575981 aaaaaacagg cgtcagttat gctgtacatt gctgcaacgg tgtacaggtc gaccgagcca
2576041 acggaatccg aataacatcc gaataatatc cttcttaccg gcgctgccat cgccattctc end of signal
2576101 atcgaagacc ctgctgacga aaaaggagca gaa**tgg**accg gaaaaccg**tg** **g**ctgggcgaa exon 2 **Trp codons**
2576161 **tgg**gtctctg tacccgagca ggacaaaaac ctggcacagt tcaagaggaa gcttcgtaag
2576221 ttttacgtct cctgtgtgac tttgatcttt ctgtacgatt gatgacgttc ttcccgaata
2576281 tatatcgttc acagagctgc ct**atg**agcca tccggaagtc aatctcaact ctactgtctt **SAHS9** “start”
2576341 ggtcaaccac ctcaagaagg gagatgaata ccatcacaag attatcatca aagaatatta exon 3
2576401 caccaatcac gtaagtttat aaccgttagc gcgctcttaa aaaaagcaaa atggtcacgc
2576461 gaaaacgttt ttcaggtcgt ttacaagctg ggcgagcagt cacccggctc gtacgacggt
2576521 ttgtcctata gtgtaaagta tggagagaaa gatggcgcgt tggttggaac ggcccattac exon 4
2576581 acaggcacca aagaccagcg tctcaacata accatgcaca acgtctacaa gctcgaaggg
2576641 gatcgtcttc tcaagagctc caccatcgac ggagtaacac tgaattgcca tcacaagagg
2576701 cgcatct**tga**a gctgtgcacg tcgtttcagt tcttcgactt ttatcgatat tcttatagtc
2576761 ttgatcaacg cttgagtaaa ggtgtttcaa gatgcataaa gactttcttc gtccttggcg
2576821 aagcttatca gtcatcttct cgaatcaaga tatccattcc atcggcagaa ggagagattc
2576881 tccgctttcc acacttgcgt gggtagactc gtgtagagcc gcgcgtatcg cggtgaacat
2576941 cgagaaaggt tcgaagcttg gcttagagtt ctcagacggt cacgggacaa gatcagatac
2577001 ggcgcaaggg tgttttcccc agcatgcaga ccgttatgac acatagttga ctttgatgcg
2577061 actggtggta ctgggaagaa tgtaacgctc caatattcgc ctaaggtgta atcctaaggt
2577121 atacgtacag tagacgcaaa cagtacagac tcctccagca acggtacttc ctggccgtat TATA box
2577181 ataaaatcaa ggttcgggtt ccttccaacc tgtacctctt tacttcttgt aactttctca
2577241 cggagaatac tcgcagaacc **atg**catcgat ttgtccttgc tctcgtcgtt tttgccggta exon 1
2577301 aacttgaaag cttcttaaga gatttaccgt cttgtgtgct atgaaagagg acgagcatga Intron 1
2577361 gctaaaaggt catcgtaaga aacgtgcgtg gctatatatg tacattttct cctggctttg
2577421 caggtgctgc catcgtctgg gccgctgatg acgctgctca cgaagaaggc gtagaa**tgg**a signal sequence
2577481 ctgggaaacc g**tgg**atgggc aaa**tgg**gaat ccgacccatc gaaggacgag aacgttgagg exon 2
2577541 aattcaaaaa gaagctccgt aagttacttt gcatttgcat ctcctcgtta gttagttttt **Trp codons**
2577601 ttagtgttca gtcgacgctg gttgtctggt ggattcgtcg aattccccgg cctgatattt
2577661 tttgcatagc tttatagatt tacccgttca gatccgtaca agcgagaaac caagacacct
2577721 atgtttgttt gcagagcttc cgatgagcca ctcggaaatg aacaaaaact ccaaagtttg
2577781 gatccatcac tacaagaagg gagacgagta ccatcacaaa atcatcatca acgacgccca exon 3
2577841 ttacaaaaac gatgtaagtc cgcaaacttt cccggttaca ttgtttctta cgtctttgct
2577901 gcacagatca aagaaaatta ttttcggctt tgtagatcgt cttcaagctg ggtcaagagt
2577961 ccgccggttc gtataacggc tcatctttca gcgtgaagta cgaggacaaa gacggcgctc exon 4
2578021 tagtcggaag cgtccactac actggcacca aagaacagtc tcttgacaag accatcaaca
2578081 acgtcttcaa gctcgaaggt gaccatctgg ttaagacttc caccatcgag ggagtgacca
2578141 tgaagcgcca ctacaacaaa cgccag**tga**a gttgtcgttg cggctaaatt ttttcctttc
2578201 tgcaaattca tgcccgtttt gtcgagtctc tcctgcttcc catcgttcta aagatttttg
2578261 cagtactgag ttatcagggc tttgtttctg ttctcgttct atcctcgtat tttcttttcg
2578321 ttcaccggat acagtaaagc tgcgtttcaa agccaggttt tttatctgcc tgttggtcgg
2578381 acggattgtc ggaccaactc agatatcgat cgggctgatt gtaaacagat actacgtatt
2578441 ttctcgtact cttcgcactg gctaacgtta ggttatacgc tctaaacggt gatcgagaaa
2578501 gttttaaaag caggcacgac atactttaaa ttctcaaacg aacagtactt ttcccatcag
2578561 atgaagcata aagcgggttt tcccagcaaa gacgcttcag cccatggcca attgaccttt
2578621 ggccgaaggc tgctggctac gaaagcgctg agtaatgata agtactcgta ctctattgtt
2578681 caactaagaa acctccgaat gtaacccaaa ggagaacgac ttcgtgccaa cttgatctta
2578741 gactgcagaa taaaccagta taaaatcaaa ccttcgccat ttcacagtac aaagcacgtt TATA box
2578801 ttaaactttt ccagccactt tcttcacgta gaagtcgagc cgcaagc**atg** aagtgtatcc **SAHS8** ex.1
2578861 tagctctcgc tctgtttgcc ggtacgtcga cagacaatta aaattttggt gctttttagt
2578921 atcatagggt aatgcaagac gttgactcga cttgtgtgga tgatcacatg agcatgtgat
2578981 tcatagttga tttcattggt ctgttgcagg tgttgccgtc gtgtgggctg gcgatgatgc
2579041 cgctcatgaa gaaggagtcg ac**tgg**acctc caagcct**tgg** ttgggcaag**t** **gg**gaatctat **Trp codons**
2579101 cccggagaag gatgaaaacc tcgtggagtt tctcaagaag ctcagtacgg tgtatagctt
2579161 tctcttcctt cagtattttc tgttctggac acatatcata aacagggtcc tcgctccagt
2579221 ccccgtccac cttagtctta gaaaatttag agttttcgag ctttacttct gttcacagat
2579281 gttcccatgg accactctaa aatgaacgcc accgtcaagg tccacctcaa ccactacaag exon 3
2579341 aaaggagacg attaccacca caagatcatc gtcaaggagg ctgagtacaa gaacgatgta
2579401 agtttgaccg ctttcgatga gttgacccgt cagctagatg accccttteg gaagtctatc
2579461 ctatggggtt tccgactgac actagacacg taatatctgt ttcgattttt aggttgtctt
2579521 caagttaggc caagagtccg ctggttcgta caacggttcg tccttcaccg tgaagtacga exon 4
2579581 agataaggat ggcgcactgg tcggaaccat tcactacacc ggtacgaagg aacagagcct
2579641 cgacaagacc atcaacaacg agtacaaggt tgaaggcaat caactggtca agacctcaac
2579701 cctcgaagga gtgacacaca agagatacta caacaaacgc aac**tga**ggtt gttcttgccg
2579761 ctatatggtt gttagttcgg ccaagttttt cctattttcg catcttttgg ctttttctca
2579821 tcattcttcc agtctttatg ttcgcctgtt ctcactgtac tttgctcaag cccatttcca
2579881 gcaacaagtg ctttattcac gtcccagaac cagctttccg ctcgtcgttt acctattcgc

```
2579941 ggaagaaatc atccaacatg accaatactg cttctgagcc gataaagtca acgattctcg
2580001 tgctggattc gatttttcga tgtagatact ggacttgact ttgacttgta cggagcgaca
2580061 ccgagcgcag caggaacgca agaagaaaac caagcagata ctttaacccg cttcaaattt
2580121 agagtgggtc tttccaggct gccactgcat tgacccgtgt tttgtttcct gaggtagact
2580181 gatccctctc ctcgaggttt caataggttc gagaacggga aagcgttgta gattacggga
```

**SAHS10 analysis**

SAHS11 and 10, shown below, are adjacent in the *R. varieornatus* genome and about 5 kilobases from the SAHS, 2, 8 cluster. The SAHS10 annotated start codon lies within a putative intron corresponding to the second intron in SAHS11 and other SAHS genes. SAHS10 encodes two of the three conserved tryptophans near the N-terminus, while the first tryptophan is replaced with a structurally plausible arginine. Upstream of this coding region, at the position of the splice acceptor found in most SAHS genes, a potential splice site is mutated, reminiscent of this site in SAHS9.

```
2568721 tcgtctgcca cgtgcacgat gccaggaacg aaaaagcaag ctaggttcag agccaacctg
2568781 ttcatacaca agtgctatcc agctatcctg ctctgcagac ttttaggcta ttctgcacag
2568841 tctccatttc tgttaactgc agttagaacg cctcaattgt catgcggcga tttcttcgca SAHS11
2568901 atcttctcat atttgccagt atgaaccttc ccacacctgt aacgttatgt gtttatgatt
2568961 tttctatttt cgtcctttga atcagaagcc tgactttcca aggataatat aagtaaacga
2569021 aatgtaatta ggtggaccga agggatctag tgttttcccc gtcgcattgg cgaatggctg
2569081 ctgttttgga taattatgat tttcgacgaa tcgataagag cttggctcaa gggactgtat
2569141 tatcccacag cttgatcctt tacaggtgcg gcagtctgcc tggcagagca cgatcctggc exon 2
2569201 catgaggaag gagccgaatg gactggaaaa ccatggctgg gcaaatgggt ttccgttccc Trp codons
2569261 gagaaggacg taaacgtact aaacttcatc acagagatcg gtcagtgcca tgatgcatca
2569321 acttcagccc tcgtggtacg ctcagcggat ggcacgggac attcgaaatg gaatgaattt
2569381 tcacaggtgt cgctgcgagt catccggaac ttcctcccat cgttacggtc ctcgtcaacc exon 3
2569441 attacaaaaa gggcgacgag taccaccaga gactgcgcgt caaggaagta gctgatcttg
2569501 atgatcacga cgtaagtacg atacacggta ctcccttgac ccaggcttaa ggaaggcttc
2569561 ggaccgacgt taacgttaag gaacctgtac tgtagattgt ctacaaactg ggccaagaaa exon 4
2569621 ccaagaacgt ttttaacggt accaccttca gtgttaagta cgatgagaaa gatgacgctc
2569681 tcgtcggaca agtcatgcta ccctcgaaca acgcgactta caagaacgag ttcaaggtcg
2569741 aaggggactt ccttgtcaag gtatcgcagc ccactttgaa ttccgcctca gccattatgt
2569801 cactggtcac atctttttc gcttgttgca gacctctgac gctcatggaa ttgtccacaa exon 5
2569861 acgatattac aagagacgga actaaatttc aagctggtgc cgaggttcag tctcagattt
2569921 tgtctttgga atcaagctcg cttgcgtgtt tgagttgtcc ccagagtaaa gtcacatgtc
2569981 gttcagttgg cgtgtcgata gaggttcttg tgcttagtct ttagtcgtac aagattttcc
2570041 cgagcagacg ggtggttcag tcggtttcca ctttttccc acattatccc gcatatctct
2570101 gtatcacaga cgaccactgc cagaaaccgc gccggggcgc ttgaccagat cagatggtga
2570161 gcaaagcagg caatcataat tcatcagcca acctttgaaa ccacgttttt ttccccgag
2570221 tgtcctaatc ggcgaatgac aatgcttgag gtcgaaaaaa agtgactaca ggctactcac
2570281 aataagggca ggtcaacact atatatgcag gcctctcatc aaaccttcca ttagtttctc
2570341 tacccatcgt actttcttac atagccagct tgaacgatgc atcgatttat ccttcttctc
2570401 gcagtctttt ccggcaagat cgagctcata tcgggatatg gttccttata ctcgggagat
2570461 ggctctgtca tcggttggct ccagccgcta actttccgtg tgcgacgtca ctgcgagaag
2570521 catcggcagg cagaatccgc actgacgcca taagaaagct taagcaagag gagttcggtt
2570581 gtcaagatgc tggatgggaa gctgtctgcg tgttttgttc gttgtgcatg tactcgtact
2570641 gtcatgatgg tgtttcgcgg gtgtggcctt tatctgggcc gccgaagacg ctgttcacga
2570701 agaaggcgta gaacggactg gcaaaccgtg gatgggcaaa tgggtcgccg ttcctgagaa Trp codons
2570761 ggacgaaaat cacgagggac tcaagaaaaa gctccgtgag tacattcgtg ttggcttctg SAHS10
```

```
2570821 accttcattc gtcagttttc ctcgacaatc gtcgatgtga tgtatgagcc accatgggga "start"
2570881 tgcgatctgc agatatcccc ttgagtcacc cgcatctgaa acacaacaac agagtgtggg
2570941 ttaacaccta caagaaggga gacgaatacc accacaagat tattatcaag gaagccggct
2571001 ataccaatga tgtacgtatg cgaagccatt atgattatgc aaactgccga acgctgcatg
2571061 gctttaatct ttccacacag tacgtgcgta ctgagatctc ttgcatacct gagactgatt
2571121 ttcaggttgt cttcaagctg ggtcaagagt ccgccggctc gcataacggc tcatctttca
2571181 gcttgaagta cgaagacaag gatggcgcct tggtcggcac cgtccatcgc accggcacca
2571241 aggaacagcc cctggacaag acgatcaaca acgtcttcaa gctcgagggt gaccatttag
2571301 ttatgacctc caccatcgac ggagtaacca tgaaacgcta ctacaagaca cgaacgtgaa
```

**SAHS12 analysis**

In the annotation of Scaffold 1 of the *R. varieornatus* genome, the ORF upstream of SAHS12 (RvY_02619-1) encodes a protein whose C-terminal region aligns well with the N-terminus of the mature SAHS proteins. A splice donor is present near the end of this sequence that corresponds to intron 2 of SAHS2 and other well-annotated SAHS genes.

This region thus may encode a large protein with the sequence MTQPMSFAQCSADRGKHSGTTIWTLLRIYLACQKAILRSKLRLRAPLFPTVEPNPAPIQNAAPAPSAAQRRRNFAASHAANVDLPGSVWHGETWGDQHDPPNRLAADVDNFDWRSKFWLGKWSSIPEKDQNLEAYLAVMgvdMNHPNMKKDQPVTLQTFKKGDKYHHKIVVEEAGYINDVIFRLGRETPGSYNGQQITVNYEEQGGALVGTVKYPAHNKVIHNTYEMDGQNLAKTSECEGVVHKRWYNKQQN, where the black amino acids are from RvY_02619-1, the "gvd" is glycine-valine-aspartate arising from the splice junction and segment upstream of the annotated SAHS12 start codon, and the brown amino acids are from SAHS12 as annotated in the Scaffold 1 annotation.

```
2002141 gatgagcaac aagtaatggc tcggtaccag ctaagacctg aggcaaaatc aacgcggatg
2002201 gatgcagatc tatttcagtg aacaaaggac ggaagatttc tgatgggagg aaaagtagta
2002261 cagtgtaata ataatccgag aaagaaaatc aaattctgac tttcagagtc atctgagatg start of
2002321 actcaaccga tgagtttcgc acagtgctcg gcggaccgcg gtaaacattc cggtaccacg RvY_02619
2002381 atctggacgc tgctgcgaat atacttggcc tgccaaaaag ccatccttcg atcaaaatta
2002441 aggctacgag gtaaacgtca caattcggac gggccaattt tctcgtcaca attcgggcgg
2002501 ctcgaaaagt tgatacgtta tttgctacgc tgtattgatt gatcaccgta gatcggttga
2002561 tacaacttta ggcttttta a cattgcgcat acccctcttc ctcccaccgc tgactccgca
2002621 actctttgac atctttcgat atctcaacct atcacaagag atagaagaga ggatgttcta
2002681 ccctgggaag caaaaactag gcagtttgga gcagaaacag ctgagataca gaccgcccaa
2002741 attgtgacga gaaaattggc ccgcccgaat tggacgtttt acctcgtaac gattatgtgt
2002801 tttgtcttct aatcagctaa tcaagtcatt ggttttcatg aggattatgt tgatgcatga
2002861 ttcctcgagc ttttgacagg ccgtaccgtg cttcacggag tgtttacccg ccgaatactg
2002921 ccaatgagtc attagaagca ccaggattgg tctgttactc tgtcctattc aaatcaggcg
2002981 aagcgggaga atgacgaagc ggatactgac gcttagcctt cgcatggtct gttgtttcgg
```

```
2003041 cttgcaccgt tccatccaga ttttagggtg agcaaccgtc agcaacgtgg ccgtcctggc
2003101 aaagtctttt cctgtcctca ccttcagctc ctttatttcc cactgtggag cctaatccgg exon 2 of
2003161 cacccatcca gaacgcagct cccgctccat cggctgccca acgtcgtcgg aactttgcag RvY_02619
2003221 ccagtcatgc cgccaatgtc gacctgccag gctccgtttg gcacggggaa acctggggag
2003281 atcaacatga tccgccgaac cgattagcag ctgatgtcga caacttcgac tggagatcga Trp codons
2003341 aattctggct gggcaagtgg agctctatcc cagagaagga tcaaaatttg gaggcttacc
2003401 ttgctgtcat gggtaagccg gagacttacg cttgattgac tgtcattgat tgactggctt
2003461 ccaacttccg cttccgggtt caacaggtgt cgacatgaac catcccaaca tgaagaagga SAHS12
2003521 tcaacccgtt acacttcaga ccttcaagaa gggtgacaag taccatcata agatcgtggt “start”
2003581 cgaggaagcc ggctacatta acgatgtaag tttgatggac ctggcagctt tcttccagcc
2003641 ggatgtcatg ttgctttatc cgacgcgaat gtagtaacgc ctttcatatt actccatgtt
2003701 gtgccgtaat ggagaagcgt ggcgataaca tgcgtgtgat acgcatggta ttccggacag
2003761 ggaggcctat gtgtcctttt attgttattc caggttattt tccgcctcgg ccgagagact
2003821 cccggatctt ataacggtca acagatcact gtcaactatg aggaacaagg cggtgctttg
2003881 gtgggtaccg tcaagtatcc cgcccataac aaggtcatcc ataataccta cgagatggat
2003941 gggcagaatc tggccaaggt atcaaacctt acttcctctt ttgcagcttt tttcctggaa
2004001 acgccggtct gacaatttgc tgacagcggt gctcgtttgt tgcagacttc cgaatgtgag
2004061 ggtgtcgttc acaagcgctg gtataacaag cagcaaaact gaagcctgtc gcctccatta
2004121 attgtgatag ttttgccttc gagttacgat tcctcatgaa agtgcttttc atgtatgtct
2004181 gccattttaa ctaactgtac cagatgttga tttacggttt tggatagctg cagtattcct
2004241 tcagagaact ttgcgatgca acgaaccatg ttccttcttt gtccactgtg aatacgatgt
2004301 gctgcgatct actatggaag cactgcctac gtagagaaaa ccgaaaatgt cctgcctcag
2004361 aactagtttc cagtttccta gacatttcga caccctccag tatctttctc gcttagggtg
2004421 tcgcacgaac agaaactaca gttctaactg cgcctttcgg cggctgactt cgcattcgaa
```
